# Extreme-Value Analysis of Intracellular Cargo Transport by Motor Proteins

**DOI:** 10.1101/2021.12.29.474400

**Authors:** Takuma Naoi, Yuki Kagawa, Kimiko Nagino, Shinsuke Niwa, Kumiko Hayashi

## Abstract

Extreme-value analysis (EVA) deals with deviations in the data from the median of the probability distributions. EVA serves various purposes such as predicting disasters and analyzing sports records. Herein, we extended the use of EVA to investigate the motility functions of nanoscale motor proteins in neurons of the living worm *Caenorhabditis elegans* (*C. elegans*). Motor proteins, such as kinesin and dynein, move along microtubules anterogradely and retrogradely, respectively, to deliver the cargo-containing materials needed for the cells. Although the essential difference between the two motors could not be inferred from the mean velocity values, the return-level EVA plots obtained from the velocity data revealed a difference. Shape parameters of the generalized extreme value distribution of EVA *ξ* < 0 for anterograde transport and *ξ* ≥ 0 for retrograde transport. Our findings extend the possibility and applicability of EVA for analyzing motility data of nanoscale proteins *in vivo*.

## Introduction

Extreme-value analysis (EVA) ^1,2^ is a statistical tool that can retrieve information regarding the extreme values of observed data that deviate from the median of probability distributions. Extreme values are important in a various topics, such as disaster prevention ^3,4^, finance ^5^, safety estimation^6^, sports ^7,8^, human lifespan ^9,10^, and the recent pandemic^11^. Recently, its applications in the biological data analysis has also become active ^12^.

‘Motor protein’ is a general term for proteins that move and function using energy obtained from adenosine triphosphate (ATP) hydrolysis; these are elaborate nanosized molecular machines that function in our bodies. For example, while myosin swings its lever arm to cause muscle contraction ^13,14^, kinesin and dynein walk along the microtubules to transport intracellular materials ^15,16^, and a part of F_o_F_1_ synthase, F_1_ rotates by hydrolyzing ATP molecules ^17^. The physical properties of motor proteins, such as force and velocity, have been investigated by *in-vitro* single-molecule experiments, in which the functions of motor proteins consisting of minimal complexes were analyzed in glass chambers ^18,19,20,21,22,23,24^. The mechanisms underlying the chemo-mechano coupling of motor proteins have been clarified by manipulating single molecules using optical tweezers ^18,19,20,21,22,23,24^, magnetic tweezers ^25,26^, and electric fields ^27,28^. Based on *in-vitro* single-molecule studies using optical tweezers, the different force-velocity relationships for kinesin and dynein have been clarified. The difference in the convexity of force-velocity relationships reflects the different mechanisms underlying the walking behavior of motor proteins on microtubules. The force-velocity relationship for kinesin is concave-down ^18^, while concave-down force-velocity relationships were found for dynein. However, its convexity remains controversial depending on the species ^19,20,21,22,24,29^.

Because motor proteins function fully in the intracellular environment and are equipped with accessory proteins, the investigation of motor proteins *in vivo* is as important as *in-vitro* single-molecule experiments; however, it is difficult to manipulate motors using optical tweezers in complex intracellular environments. Without any manipulation, we found that EVA successfully provided information regarding the convexity of the force-velocity of motor proteins *in vivo*.

Here, we extended the use of EVA to investigate nanoscale phenomena associated with the function of motor proteins inside cells, focusing on the *in vivo* velocity of synaptic cargo transport performed by the motor proteins kinesin (UNC-104 ^30,31^) and dynein ^32^ in the axons of motor neurons of living *Caenorhabditis elegans* (*C. elegans*), a model organism in neuroscience. As the axons of these worms are sufficiently long, this *in vivo* system is appropriate for investigating intracellular cargo transport. Synaptic materials packed as cargo are delivered to the synaptic region of neurons via kinesin-mediated anterograde transport, and unnecessary materials that accumulate in the synaptic region are returned to the cell body via dynein-mediated retrograde transport (Fig. 1a). Since the bodies of worms are transparent and their body movement is suppressed by anesthesia, the motion of fluorescently labeled synaptic cargos in living worms can be observed by fluorescence microscopy (Fig. 1b) ^33^. Velocities were measured from recorded images. By applying EVA to the velocity data of the intracellular cargo transport, we investigated the force-velocity relationship between kinesin and dynein. We found that difference in the parameters of the generalized extreme value distributions and return level plots of the EVA. The difference was then related to the force-velocity relationship of kinesin and dynein by using simulations.

**Fig. 1.**
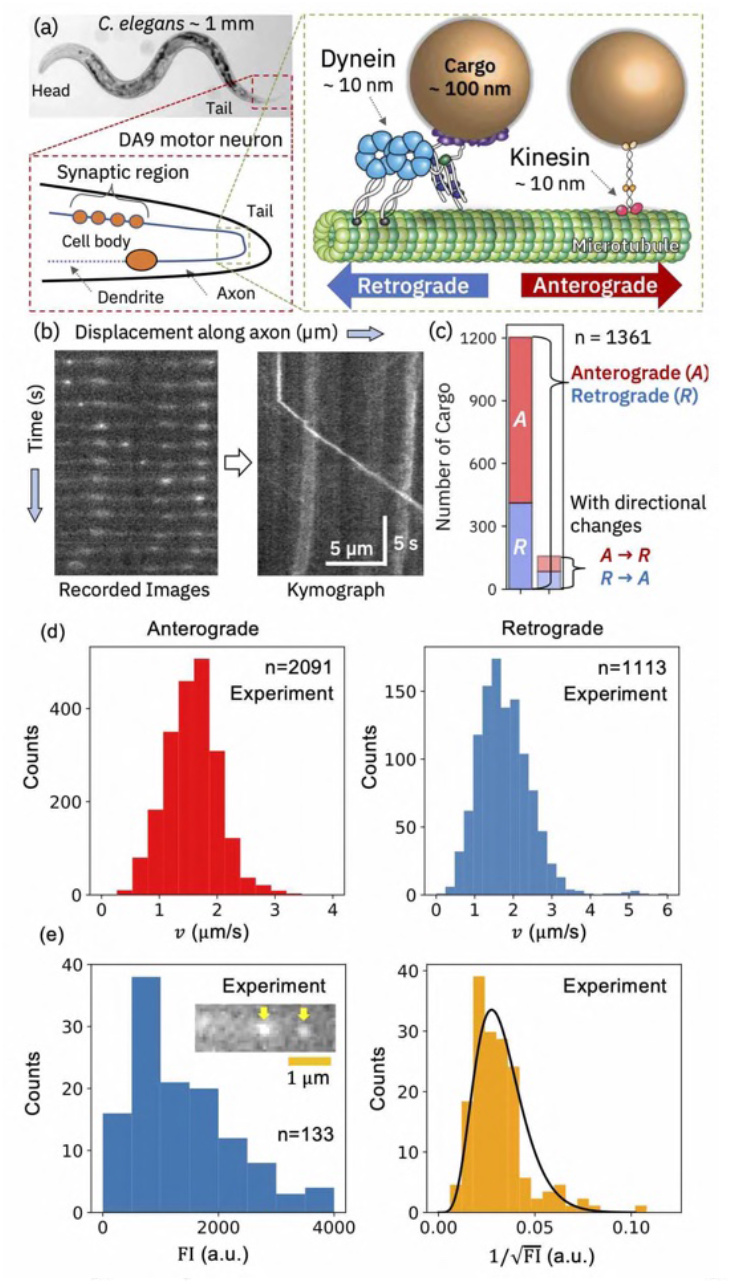
(a) Schematics of anterograde and retrograde synaptic cargo transport by kinesin and dynein, respectively, in the DA9 motor neurons of *C. elegans*. (b) Kymograph analysis. Velocity (*ν*) was measured as the slope of the trajectory of a fluorescently labeled cargo. (c) Number of synaptic cargos moving anterogradely (A), retrogradely (R), and exhibiting direction change (A→R and R→A). (d) Histogram of the velocity {ν^*i*^} of synaptic cargos for anterograde (red) and retrograde (blue) transport. (e) Fluorescence micrographs of synaptic cargos (inset). Fluorescence intensity (FI) of synaptic cargos (*n* = 133) (left panel). The distribution of 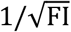 was fitted using a Gamma distribution *b*^*a*^*x*^*a-1*^*e*^*-bx*^/Γ(*a*) with *a* = 6.5, *b*= 0.0050 (right panel).

## Results

### Transport velocity of synaptic cargos

Fluorescence images of green fluorescence protein (GFP)-labeled synaptic cargo transported by motor proteins were captured using a 150× objective lens and an sCMOS camera at 10 frames per second (see Methods). The body movements of *C. elegans* worms were suppressed under anesthesia. For only the motionless worms, kymograph analysis of the recorded images was performed using the ‘multi kymograph’ module in ImageJ ^34^ (Fig. 1b). Although we observed 562 worms totally, the velocity values of the moving cargo were collected from 232 worms, revealing that transport velocities were not observed in most of the worms because of the body movement and the obscurity of fluorescence movies.

The velocity values were calculated as the slopes of the trajectories of the fluorescently labeled cargo in the kymograph images. Typically, a cargo exhibits a moving motion at a constant velocity, and pauses, and rarely reverses its direction (Fig. 1c). Constant velocity segments (CVSs) were assumed to be unaffected by the other motor proteins much based on other experimental studies ^35,36^. (It means that the tug-of-war between the two motors, kinesin and dynein^36,37^, was not considered for CVSs, and that we supported the “motor coordination model,” in which adaptor proteins that connect motors with cargos deactivate opposing motors ^32^.)

Figure 1d presents the histograms of the measured velocities {*ν*^*i*^} (*i* = 1, ⋯, *n* where *n* =2091 for anterograde transport from 228 worms and 0=1113 for retrograde transport from 217 worms). The mean velocities were 1.6±0.5(SE) μm/s and 1.8±0.7(SE) μm/s for anterograde and retrograde transport, respectively. These values are similar to those reported for the same neurons in *C. elegans* worms^38,39^ (higher than the values obtained by the *in vitro* single-molecule experiments^21,39,40^). In the *in vivo* case, the velocity values varied widely as revealed by the histogram (Fig. 1d). This broad variation was considered to originate from the cargo size difference (Fig. 1e) based on our previous study ^41^. Here, the functional form of distribution of the cargo size was estimated from that of the fluorescence intensity (FI) of the cargo (Fig. 1e) through the relation FI ∝ 4*πr*^2^ (*r*: radius of cargo), *i*.*e*., 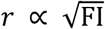, assuming that the fluorescent proteins labeling a cargo are uniformly distributed on its surface.

### Application of EVA to assess transport velocity data

In this study, one block of EVA was considered as a single worm. Approximately 5–20 velocity values were observed for each worm (a block), from which the largest value 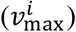 was selected. Using 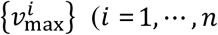 where 0=228 for anterograde transport and 0=217 for retrograde transport), the return-level plot was investigated (Figs. 3a, b). The two axes of the return level plot represent the return period *r*_*p*_ and return level *z*_*p*_. For a given probability *p, r*_*p*_ = −{log (1 − *p*) }^−1^, and *z*_*p*_ is defined by the generalized extreme value distribution as 1 − *p* = *G*(*z*_*p*_), where

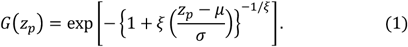

(Note that *z*_*p*_ and *r*_*p*_ represent 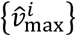 and the number of samples, respectively. Here 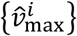 is the rearranged data of 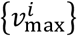, such that 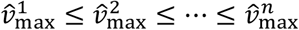 From EVA by using the ‘ismev’ and ‘evd’ packages in R ^42^, we obtained parameters of the generalized extreme value distribution *ξ, μ*, and *σ* in Eq. (1) (Supplementary Table S1). We found that *ξ* < 0 for anterograde transport and *ξ* ∼0 for retrograde transport. The return level plot for anterograde transport shows a convergent behavior as *r*_*p*_ becomes larger (Fig. 2a), a property specific to a Weibull distribution (*ξ* < 0), and *V*_max_ was proved to exist in this case and was estimated to be 4.0±0.4 μm/s using the following equation:

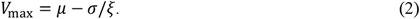

**Fig. 2.**
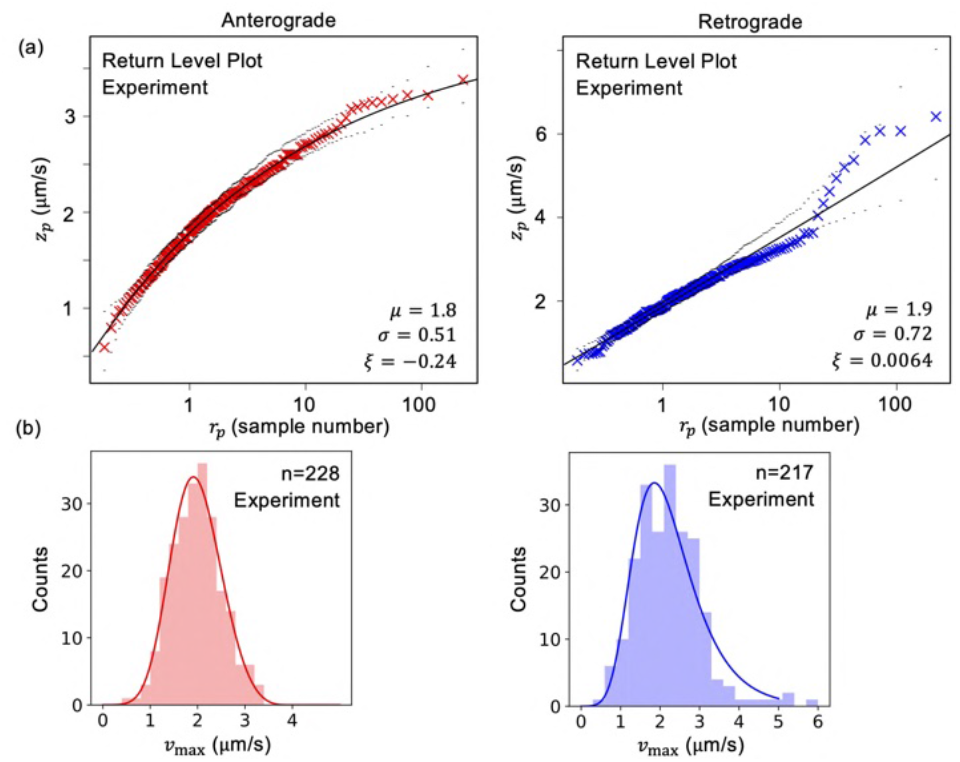
(a) Return-level plots experimentally measured for anterograde and retrograde transport in the neurons of *C. elegans* worms. The black dotted lines represent the reliable section. (b) Distributions of *ν*_max;_ for anterograde and retrograde transport.

However, the return-level plot of the retrograde transport (Fig. 2b) shows *ξ*∼ 0, and *V*_max_ cannot be estimated using Eq. (2). Figure 2b shows the distributions of 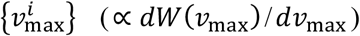 for anterograde and retrograde transport.

We checked that the results *ξ* < 0 for anterograde transport and *ξ* ∼0 for retrograde transport did not depend on the selection of samples from the bootstrapping analysis of the data 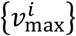 (Supplementary Fig. S1). We also investigated the block sizes (the number of worms) from which 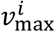 was selected, which did not affected on the result, neither (Supplementary Fig. S2).

### Construction of a simulation model using the force-velocity relationship

We considered the behaviors of the return-level plots for anterograde and retrograde transport from the viewpoint of the force-velocity relationship of motor proteins. According to the results of previous studies using single-molecule experiments, two regimes exist in the force-velocity curves of motor proteins: the load-sensitive and load-insensitive regimes ^43^ (Fig. 3a). In the load-sensitive regime, the velocity changes rapidly with an increase in load (*F*), whereas in the other regime, the velocity changes only slightly with an increase in load (*F*). *In vitro* single-molecule experiments revealed that the force-velocity curve of kinesin was concave-down ^18^, whereas some dynein data showed concave-up ^21,22,24^. This mechanical difference in the force-velocity relationship can be explained as follows: kinesin keeps moving at a distance of approximately 8 nm along a microtubule (the interval of the microtubule structural unit) per hydrolysis of a single ATP molecule even when a low load is applied, which makes its force-velocity relationship load-insensitive, resulting in a concave-down force-velocity curve. However, some dyneins, whose force-velocity relation shows a concave-up relation and variable step sizes of 8–40 nm under no-load conditions ^19,21,23,44^, slow down rapidly by decreasing the step size even when a low load is applied, resulting in a rapid velocity decrease and a concave-up force-velocity curve. Because several study results show that dynein takes an 8-nm step like kinesin^20,29^, in this case, the larger load dependence of the velocity of dynein is explained by the load-dependent kinetic rates. The velocity in the low load condition is written as *ν*(*F*) (μm/s) ∼*ν*(0) − 0.04*F* (pN) for kinesin and *ν* (*F*) (μm/s) ∼*ν*(0) − 0.2*F* (pN) for dynein from the kinetic rates of the three-state model ^45^.

**Fig. 3.**
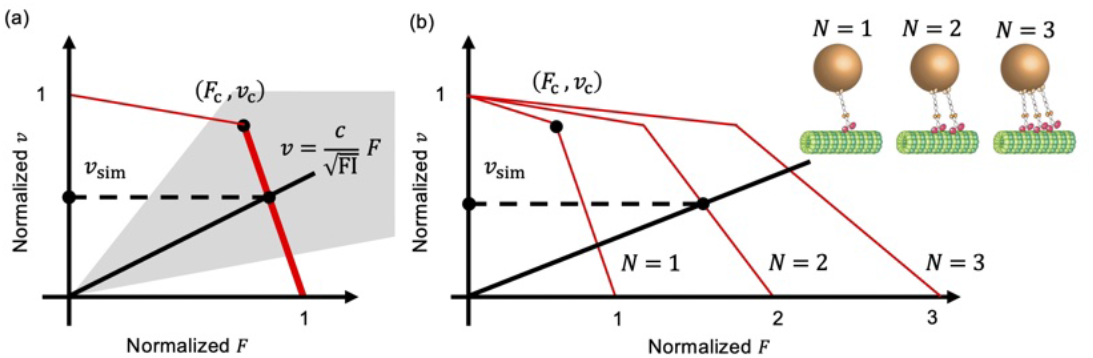
(a) Schematics of a concave-up force-velocity relationship of one motor protein. Two regimes, load-insensitive (thin colored line) and load-sensitive (thick colored line) regimes, are represented. The thick black line represents the Stokes ’ law 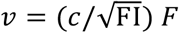. The grey area represents the possible values of 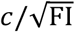, decided from the experimental velocity values. *N*orm*a*liz*e*d *F* : *F*/*F*S and ν : *ν*/*V*_m*ax*_,where *F*_S_ is the stall force of a motor protein. (b) The case of the force-velocity relationship of multiple motors (*N* = 3).

The simplest model of the force-velocity curve can be characterized by the changing point (*F*_c_, *ν*_c_) (Fig. 3a) between the load-sensitive and load-insensitive regimes. In the following sections, we aim to find the region (*F*_c_, *ν*_c_) that reproduce the return-level plots shown in Figs. 2a and b by performing numerical simulations. Note that in the following simulation, the axes of the force-velocity relationship are normalized as (*F*/*F*_s_, ν/*V*_max_), where *F*_s_ is the stall force of a motor protein ^18,19,20,21,22,23,24^, which is the maximum force generated by the motor against an opposing load. We used (*F*_s_, *V*_max_) = (8 pN, 4 μm/s) for anterograde transport and (*F*_s_, *V*_max_) = (7 pN, 6.5 μm/s) for retrograde transport. The *F*_s_ values were obtained from the reference ^45^. *V*_max_ for anterograde transport was determined from the maximum values using Eq. (2), and that for retrograde transport was the maximum of the observed experimental velocities as an approximate value of *V*_max_ because *V*_max_ could not be estimated using Eq. (2).

From a given (*F*_c_, *ν*_c_), the simulated velocity value *ν*_sim_ is obtained as the intersection between the black line 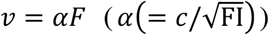 representing the Stokes’ law and the force-velocity curve *ν* = *f*(*F*) (*F* = *f*^−1^(*ν*)) of a motor protein (Fig. 3a). The value of 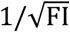 is stochastically generated from the Gamma distribution 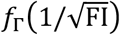 (Fig. 1e, right), noting that 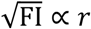 where FI and *r* are the FI and radius of a cargo, respectively. The constant *c* is chosen so that the range (=50 % interval) of the Gamma distribution multiplied by *c* corresponds to the range of the measured velocity distribution: 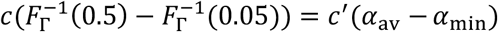. *F*_Г_ is the cumulative distribution of 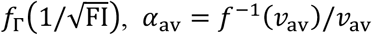 and *α*_min_ = *f*^−1^ (*ν*_min_)/ *ν*_min_ where *ν*_av_ and *ν*_min_ correspond to the mean value and minimum of the measured velocities. *c*^′^ is a tuning parameter so that the variance of the experimentally measured velocity distribution is similar to the variance of *ν*_sim_. (To summarize, we decided *α* in *ν* = *αF* model so that the cargo distribution in Fig. 1e represented the variations of the measured velocities (Fig. 1d) approximately.) This procedure was repeated 2000 times (*i*.*e*., *i* = 1, ⋯, *n* where *n*=2000) (Fig. 4a). Subsequently, 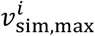 was chosen from among the 10 values of *ν*_sim_.

**Fig. 4.**
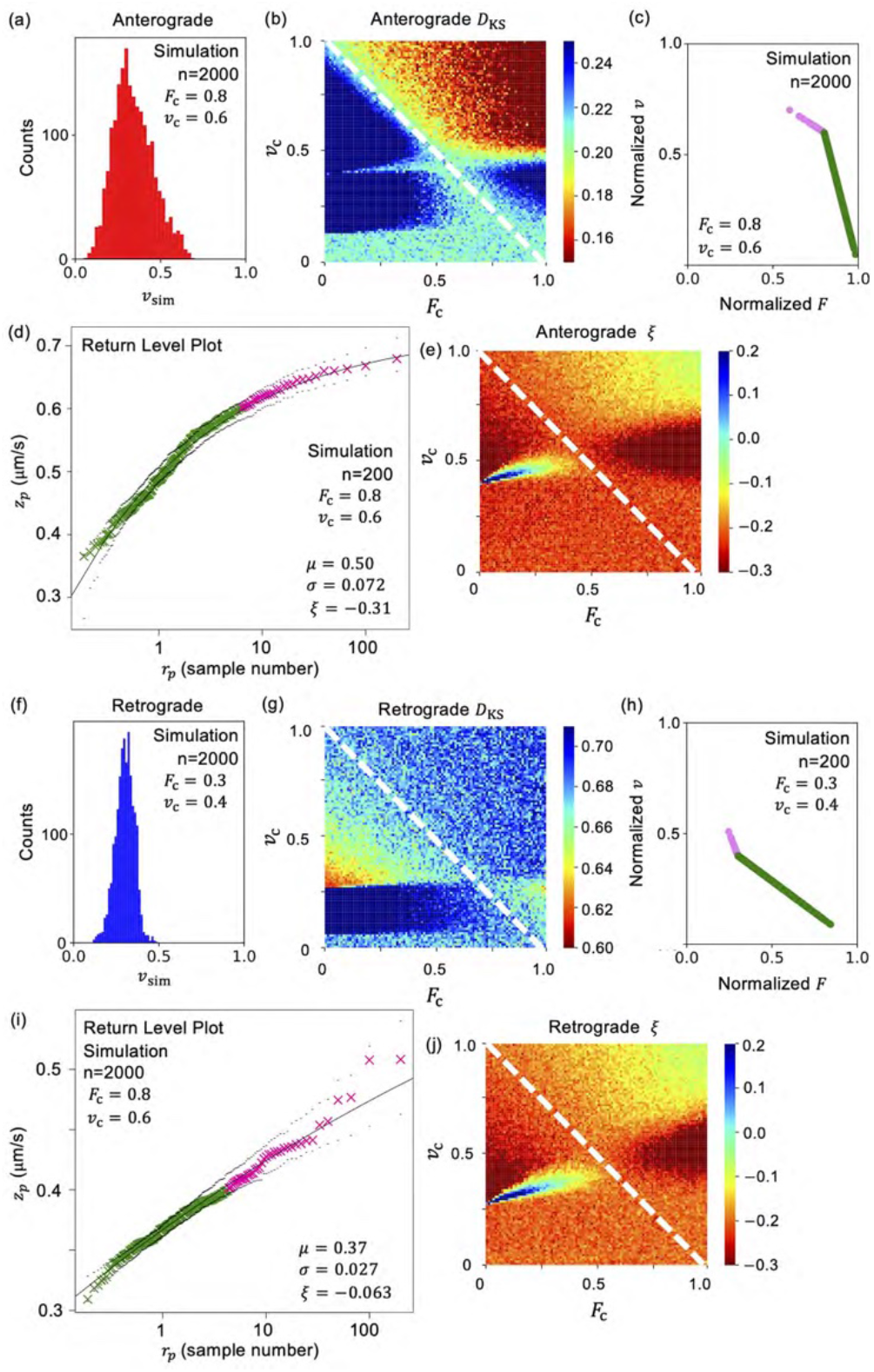
Simulation results in the case depicted in Fig. 3a for anterograde (a–e) and retrograde (f–j) transport. (a,f) Distribution of *ν*_Sim_. (b,g) Contours of *D*_kS_. The value of *D*_kS_ is the average of five trials. (c,h) (*ν*_Sim_, *f* -1(*ν*Sim)) for (*F*_c_, *ν*_c_). (d,i) Return-level plots of 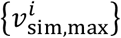 (e,j) Contours of shape parameter *ξ*. The value of *ξ* is the average of five trials.

### Outline of force-velocity relationships decided by the comparison between the results obtained from the experiment and simulation

The Kolmogorov-Smirnov statistic D_KS_ is measured for the dataset 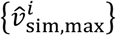

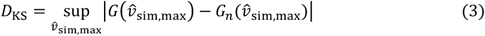

Here, the extreme-value dataset 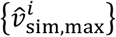 is the rearranged data of 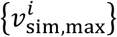, such that 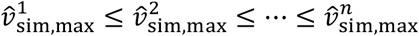, and 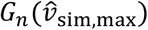 is defined as (the number of elements in the sample 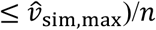. The set of the parameters (*ξ, μ, σ*) for measured values (Fig. 2 and Supplementary Table S1) are used to construct *G* (Eq. (1)). *D*_KS_ was calculated as the mean value of five trials for each (*F*_c_, *ν*_c_).

In the case of anterograde transport, Fig. 4b represents the contour of *D*_KS_. *D*_KS_ was smaller when (*F*_c_, *ν*_c_) is in the region above the diagonal line (white) of the graph. For the typical case of the region (*F*_c_, *ν*_c_) = (0.8,0.6), Figs. 4c and 4d show the force-velocity relationship 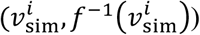 and the return level plot, respectively (for 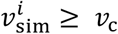, the symbols are plotted as the pink circles). Because 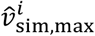 for a large *r*_*p*_ (pink symbols) was chosen from the load-insensitive regime of the force velocity relationship, it showed convergent behavior, and the graph shows a Weibull-type behavior similar to the experimental one (Fig. 2a). The calculated results (the contour) for the shape parameter of the generalized extreme value distribution, *ξ*, for each (*F*_c_, *ν*_c_) are plotted in Fig. 4e. The characteristic of the Weibull distribution *ξ* < 0 is valid for the region above the diagonal line (white) in Fig. 4e. It implied that *ξ* tends to become negative for the concave-down force-velocity relationship.

Figures 4f–j are the results of the retrograde transport case (the model parameters, *ν*_av_ and *ν*_min_, *ξ, μ, σ* are the measured retrograde ones). Fig. 4g represents the contour of *D*_KS_. In the case of retrograde transport, *D*_KS_ decreases when (*F*_c_, *ν*_c_) is chosen from the region below the diagonal line (white) of the graph. For the typical case of the region (*F*_c_, *ν*_c_) = (0.3,0.4), the force-velocity relationship 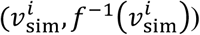 and the return level plots are shown in Figs. 4h and 4i, respectively (for 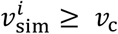, the symbols are plotted as the pink symbols). Unlike the anterograde case, 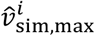 for a large *r*_*p*_ (pink symbols), belonging to the load-sensitive regime of the force velocity relationship, easily created a gap. Because a small cargo size, which generates a large 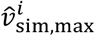, is a rare event based on the FI distribution (Fig. 1e), the gaps easily generated and 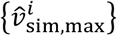 was hard to converge in the return level plot in the load-sensitive regime. We discussed the effect of cargo size distributions in the Discussion section (see also Supplementary Fig. S4).

The calculated results for the shape parameter of the generalized extreme value distribution, *ξ*, are plotted in Fig. 4j. *ξ*∼0 or *ξ* > 0 is typically observed for the blue region of (*F*_c_, *ν*_c_). This implies that *ξ* tends not to be negative for the concave-up force-velocity relationship. Note that the simulation results for 0 = 400 and 0 = 1000 are shown in Supplementary Fig. S3, to check that the gaps did not vanish as the number of sample 0 becomes large.

### Effects of cooperative transport by multiple motors

It has been suggested that a single cargo can be transported using multiple motors (Fig. 3b). The effects of multiple motor transports on the results of the present study were investigated (Fig. 5). Previously, we used a non-invasive force measurement technique ^41,46,47^ developed by our research group to examine cargo transport in the neurons of *C. elegans*, and estimated that the number of motors carrying synaptic cargo was 1–3 ^33,48^. The frequency *P*(*N*) of the number of motors (*N* = 1,2,3) carrying the cargo was approximately *P* (1): *P* (2): *P* (3) = 1: 2: 1, based on the previous observation ^33^. After selecting *N* according to *P* (*N*), *ν* (*F* / *N*) was used instead of *ν* (*F*) to determine *ν*_sim_ from the intersection between *ν* (*F* / *N*) and the line *ν*= *αF* (Fig. 3b). The EVA was applied to {*ν*_sim_} using the same procedure as that used to obtain the results depicted in Fig. 4. The variance in the velocity distribution increases in the case of multiple motor transport (Figs. 5a and 5f), which is similar to the experiments. The other tendencies of the contours of D_ks_ and *ξ* are similar to the case of one motor (Fig. 4).

**Fig. 5.**
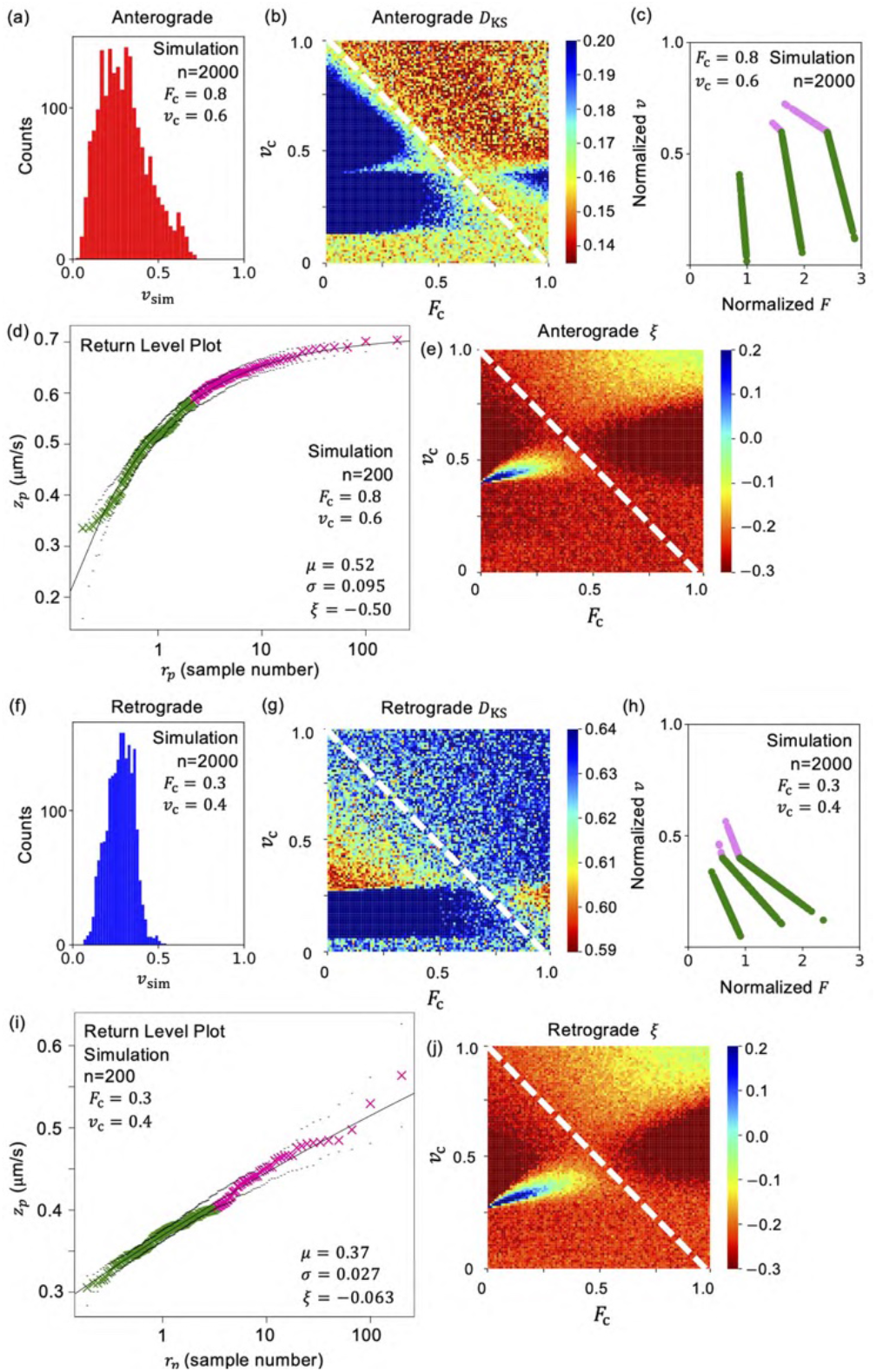
Simulation results in the case depicted in Fig. 3b (multiple motor case) for anterograde (a–e) and retrograde (f–j) transport. (a,f) Distribution of *ν*_*S*im_. (b,g) Contours of *D*_kS_. The value of *D*_kS_ is the average of five trials. (c,h) (*ν*_Sim_, *f*^-1^(*ν*_Sim_)) for (*F*_c_, *ν*_c_). (d,i) Return-level plots of 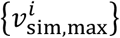 (e,j) Contours of shape parameter *ξ*. The value of *ξ* is the average of five trials.

### Chemo-mechanical coupling models of the force-velocity relationship

We referred to the force-velocity relationship between kinesin and dynein, which is theoretically derived based on the mechanisms underlying ATP hydrolysis by motor proteins. The force-velocity relationships were derived from the three-state model ^45^. The force-velocity relationship for the three-state model of ATP hydrolysis is represented as follows:

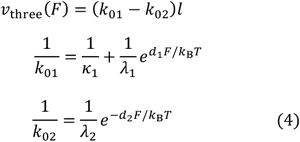

See reference^45^ for the definitions of parameters for both anterograde and retrograde transport. Subsequently, the simple force-velocity relationship *v*(*F*) depicted in Fig. 3a was replaced with this *v*_three_(*F*). The differences in the model parameters (Eq. (4)) was resulted in the different convexities of the force-velocity relationship (Fig. 6a). In Fig. 6a, the circles represent the 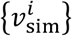 values obtained from the simulations. 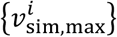 were chosen from 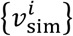. Using these 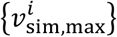, we calculated the return-level plots for the three-state model in for *n =* 200, 400 and 1000. We found that the tendency that *z*_*p*_ did not converge for a large *r*_*p*_ in the case of the retrograde transport, *i*.*e*., the gaps created between 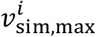 and 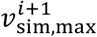 for a large *i*. This is because a large velocity value is likely to be generated in the case of a concave-up force-velocity relationship, owing to its steep slope in the load-sensitive regime.

**Fig. 6.**
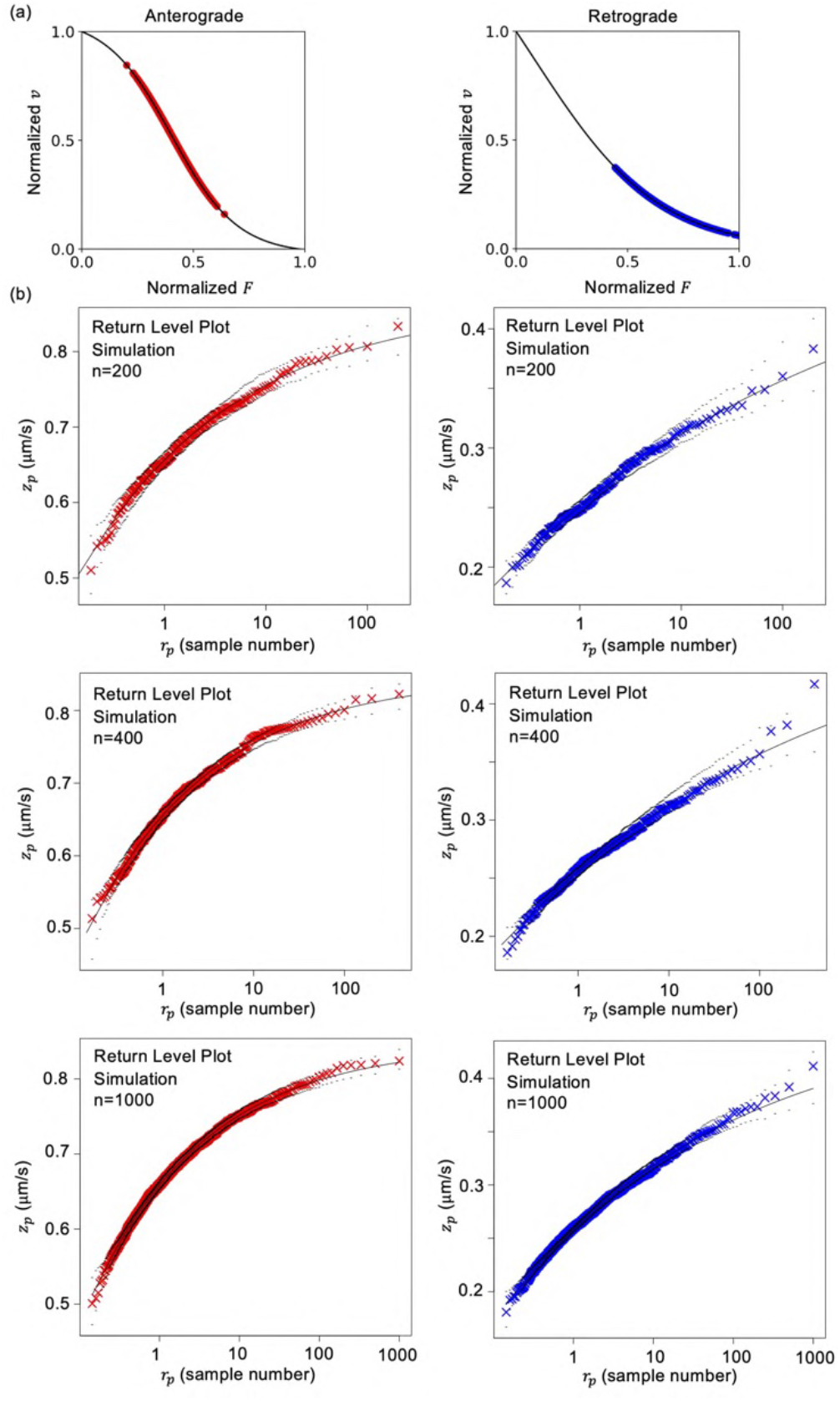
Force-velocity relationship (a) and corresponding return-level plots (b) of the three-state model (Eq. (4)) in the cases *n* = 200, 400 and 1000 for anterograde and retrograde transport.

## Discussion

To the best of our knowledge, we applied EVA, for the first time, to assess cargo transport by the motor proteins kinesin and dynein in the neurons of living worms, as observed by high-resolution fluorescence microscopy. We investigated the velocities of the transport and found that the return-level plots of the extreme velocity values revealed differences between the motor protein types. The return level plot of anterograde transport by kinesin shows the typical behavior of a Weibull distribution (the shape parameter *ξ* < 0), where the Weibull type data has a maximum value (Eq. (2)), the counterpart of retrograde transport by dynein does not show a Weibull type behavior (non-negative shape parameter *ξ* ∼0 or *ξ* > 0). Using the simulation, the abnormality of the return-level plot that appeared only for the retrograde velocity data was attributed to the fact that the force-velocity relationship for retrograde transport was concave-up, whereas that for its anterograde counterpart was concave-down. The steep velocity decrease in the low-load condition for the retrograde transport caused a major variation in the larger velocity values, and this behavior tends to generate a major variation in velocity near the maximum velocity and *ξ* > 0 as a result. This phenomenon was occurred because the appearance of large velocity values in the case of small cargo sizes (small values of 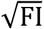) was a rare event. When a small cargo size is not a rare event–for example, when the cargo size distribution is uniform unlike the case of the distribution in Fig. 1e–the retrograde velocity data showed a Weibull type in the simulation (Supplementary Fig. S4), and had a maximum value based on Eq. (2). No large gaps generated between 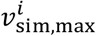 and 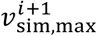even for a large *i* in this case.

Recent *in vitro* single-molecule experiments have suggested a concave-up force-velocity relationship for ciliary ^22^ and mammalian dynein ^21,44^, whereas yeast dynein exhibits a concave-down (kinesin-like) force-velocity relationship ^19,20^. In the present study, we found a concave-up force-velocity relationship for cytoplasmic dynein in *C. elegans*. To investigate mammalian dynein, EVA was also applied to examine synaptic cargo transport in mouse hippocampal neurons, as originally reported in a previous study ^41^. The return-level plot with *ξ* > 0 was also observed for retrograde transport (Supplementary Fig. S5), which corresponds to the concave-up force-velocity relationship reported in previous studies ^21,44^.

Interestingly, several dynein motors exhibit a concave-up force velocity curve. The biological significance of collective cargo transport by multiple motor proteins is explained below and was first introduced in a previous study ^44^. When multiple motors work together, the leading dynein decreases its velocity rapidly in the presence of a low load so that the trailing dynein can catch up. This allows the trailing dynein to share the load with the leading dynein, thereby preventing detachment of the leading dynein from the microtubules. In other words, the rapid decrease in velocity in the load-sensitive region results in the self-correction of the position of dynein molecules, allowing them to move as a loosely bunched group ^44^. However, the leading kinesin does not slow with regard to the concave-down force-velocity relationship in the presence of a low load. Consequently, the trailing kinesin cannot catch up with the leading kinesin, causing it to easily detach from the microtubules ^44^. (Because the load acting on each motor may be different in such multiple-motor cases, there is room to improve the equal load share model between the motors (*F = F*_total_/*N*) used in our simulation (Fig. 3b) in reference to previous models ^49,50,51,52^.)

Although the outlines of the *in vivo* force-velocity relationships could be inferred using the EVA, the stall force values regarding the maximum forces of the motors could not be estimated from this analysis. Many *in vitro* single-molecule studies have provided the stall force values of kinesin and dynein using optical tweezers ^18,19,20,21,22,23,24^. Several challenging attempts have made for *in-vivo* force measurement ^53,54,55^.

Interpretation of return-level plots based on the *in vivo* force-velocity relationship is a promising tool for future research regarding neuronal diseases, particularly, KIF1A-associated neurological disorders^39,56,57^. KIF1A is a type of kinesin transporting synaptic vesicle precursor cargos, and the force and velocity of the pathogenic mutant KIF1A have been reported to be impaired ^56,57^. Since *in vivo* force measurement is difficult, the estimation of *in vivo* physical properties using EVA can be helpful for understanding the *in vivo* behavior of motor proteins. Thus, we believe that the findings of the present study represent a step forward toward broadening the scope of EVA applications.

## Methods

### Sample preparation

In our study, we used *C. elegans* stains *wyIs251*[Pmig-13::gfp::rab-3; Podr-1::gfp]; *wyIs251* has been previously described ^58,59^.

### Culture

*C. elegans* was maintained on OP50 feeder bacteria on nematode agar plates (NGM) agar plates, as per the standard protocol ^58,59^. The strains were maintained at 20°C. All animal experiments complied with the protocols approved by the Institutional Animal Care and Use Committee of Tohoku University (2018EngLMO-008-01, 2018EngLMO-008-02).

### Fluorescence microscopy observations

A cover glass (32 mm × 24 mm, Matsunami Glass Ind., Ltd., Tokyo, Japan) was coated with 10% agar (Wako, Osaka, Japan). A volume of 20 μL of 25 mM levamisole mixed with 5 μL 200-nm-sized polystyrene beads (Polysciences Inc., Warrington, PA, USA) was dropped onto the cover glass. The polystyrene beads increased the friction and inhibited the movement of worms; levamisole paralyzed the worms. Ten to twenty worms were transferred from the culture dish to the medium on the cover glass. A second cover glass was placed over the first cover glass forming a chamber, thereby confining the worms. The worms in the chamber were observed under a fluorescence microscope (IX83, Olympus, Tokyo, Japan) at room temperature. Images of a GFP (green fluorescence protein)-labelled synaptic cargos in the DA9 motor neuron were obtained using a 150× objective lens (UApoN 150x/1.45, Olympus) and an sCMOS camera (OLCA-Flash4.0 V2, Hamamatsu Photonics, Hamamatsu, Japan) at 10 frames per second.

### Error of *V*_max_

The mean 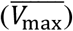 and error (*δV*_max_) of *V*_max_ were estimated from the fitting parameters 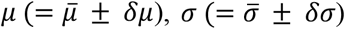 and 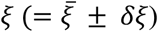 of the Weibull distributions defined by Eq. 1 in the main text as follows:

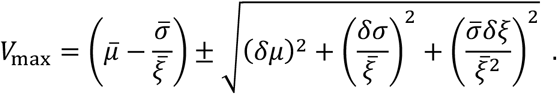

### Bootstrapping method

A part (*r*: ratio) of the data 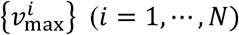 was randomly selected, and then the fitting parameters of *μ, σ* and *ξ* in Eq. (1) were calculated for the partial data. (Here, duplication of the same data was allowed if it was selected.) This procedure was repeated 10 times, and then the errors of *μ, σ* and *ξ* were calculated. In Supplementary Fig. S1, each parameter is plotted as a function of *r* in the cases of anterograde and retrograde transport. The values were stable for a wide range of 6 (0.6 ≤ 6 ≤ 1).

### Block size

We investigated the dependence of the fitting parameters (μ, σ and *ξ*) on the number (s) of worms (block size), from which 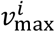 was chosen, as shown in Supplementary Fig. S2.

## Acknowledgments

We acknowledge Dr. T. Saigo for facilitating useful discussions regarding EVA and Dr. K. Sasaki for facilitating useful discussions regarding motor proteins. We would like to thank Edita<u>g</u>e (www.editage.com) for their assistance with English language editing. This work was supported by the JST PRESTO (Grant No. JPMJPR1877), and the FRIS Creative Interdisciplinary Research Program, Tohoku University to K. H., and by JSPS KAKENHI (grant nos. 19H04738, 20H03247, and 20K21378) to S. N.

## Author contributions

K.H. conceived the project with the help of S.N and wrote the paper. T.N. analyzed the data and performed the simulation. Y.K. and K.N. performed the experiments. S.N. provided the sample.

## Data availability

Data supporting the findings of this study are available within the article and its Supplementary Information files, and also from the corresponding author on reasonable request.

## Supplementary Information

**Table S1.** Values of the fitting parameters *μ, σ* and *ξ* (experiments of *C. elegans* and mouse). The errors represent the fitting errors estimated by the maximum likelihood method (MLE).

| *C. elegans* | Anterograde | Retrograde |
| --- | --- | --- |
| $\mu$ ($\mu\text{m/s}$) | $1.79 \pm 0.04$ | $1.86 \pm 0.054$ |
| $\sigma$ ($\mu\text{m/s}$) | $0.508 \pm 0.026$ | $0.716 \pm 0.038$ |
| $\xi$ | $-0.235 \pm 0.042$ | $0.00643 \pm 0.04015$ |
| $V_{\text{max}}$ ($\mu\text{m/s}$) | $3.95 \pm 0.40$ | — |

**Table S1** Values of the fitting parameters $\mu$ , $\sigma$ and $\xi$ (experiments of *C. elegans* and mouse). The errors represent the fitting errors estimated by the maximum likelihood method (MLE).
| Mouse | Anterograde | Retrograde |
| --- | --- | --- |
| $\mu$ ($\mu\text{m/s}$) | $1.91 \pm 0.10$ | $1.30 \pm 0.07$ |
| $\sigma$ ($\mu\text{m/s}$) | $0.970 \pm 0.078$ | $0.648 \pm 0.054$ |
| $\xi$ | $-0.261 \pm 0.093$ | $0.107 \pm 0.072$ |
| $V_{\text{max}}$ ($\mu\text{m/s}$) | $5.62 \pm 1.36$ | — |

**Figure. S1.**
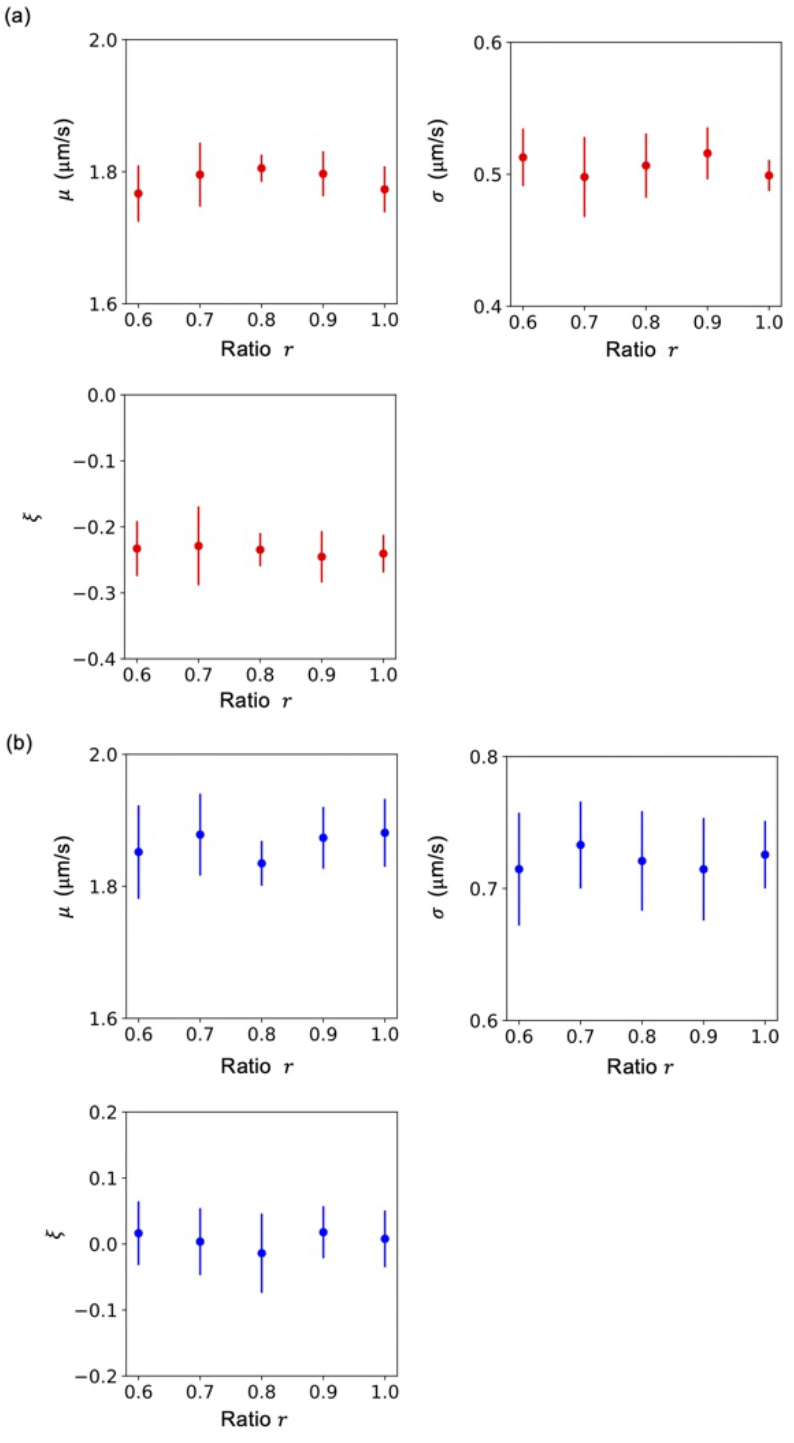
Results of the boot strapping method (*C. elegans*). Fitting parameters of *μ, σ*, and *ξ* plotted as a function of ratio (*r*). The bootstrapping analysis was repeated 10 times for each *r*. The error-bars represent the standard deviations (SD).

**Fig. S2.**
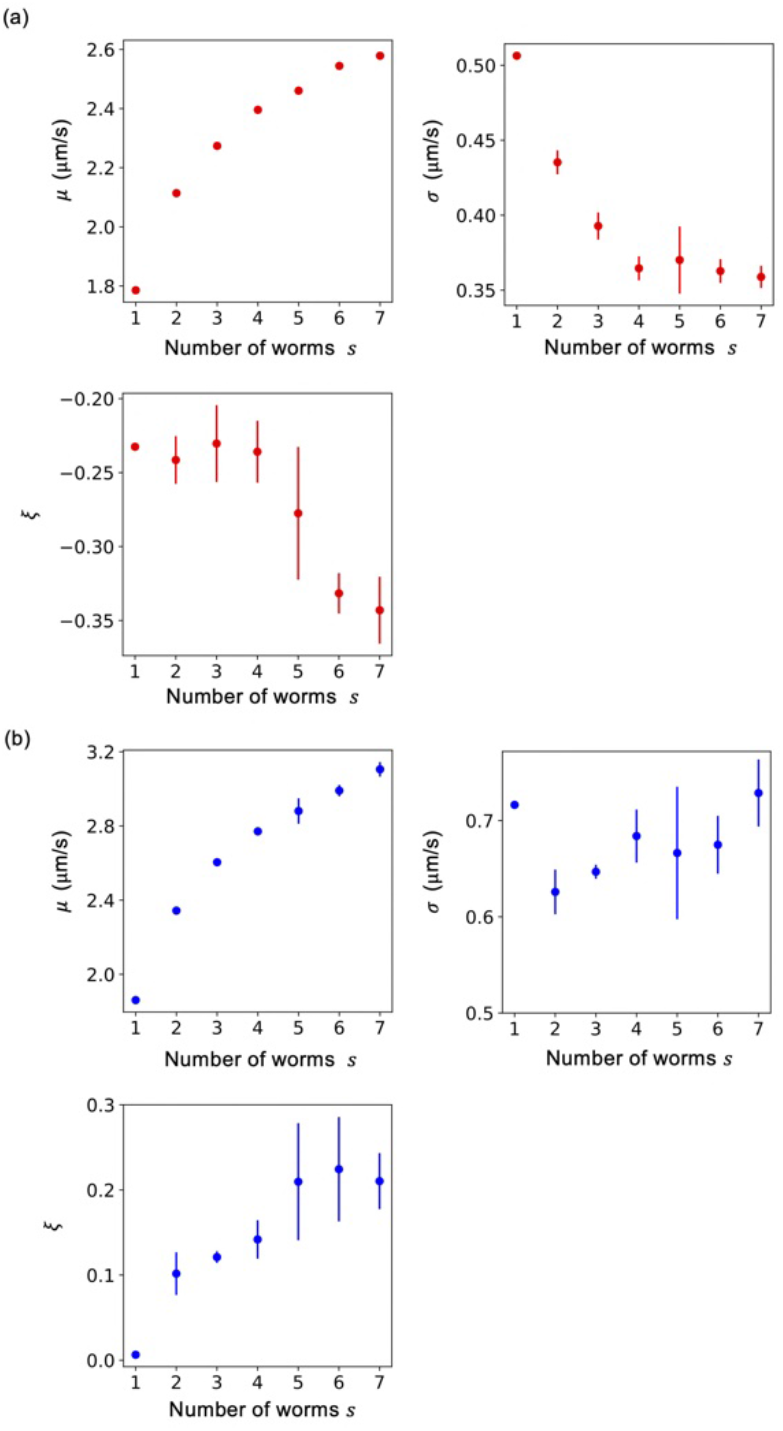
Block size dependence (*C. elegans*). Fitting parameters of *μ, σ*, and *ξ* plotted as a function of the number (*s*) of worms (block size). The error-bars represent the standard deviations (SD) for the 10 repeats of bootstrapping trials (*r*=1).

**Fig. S3.**
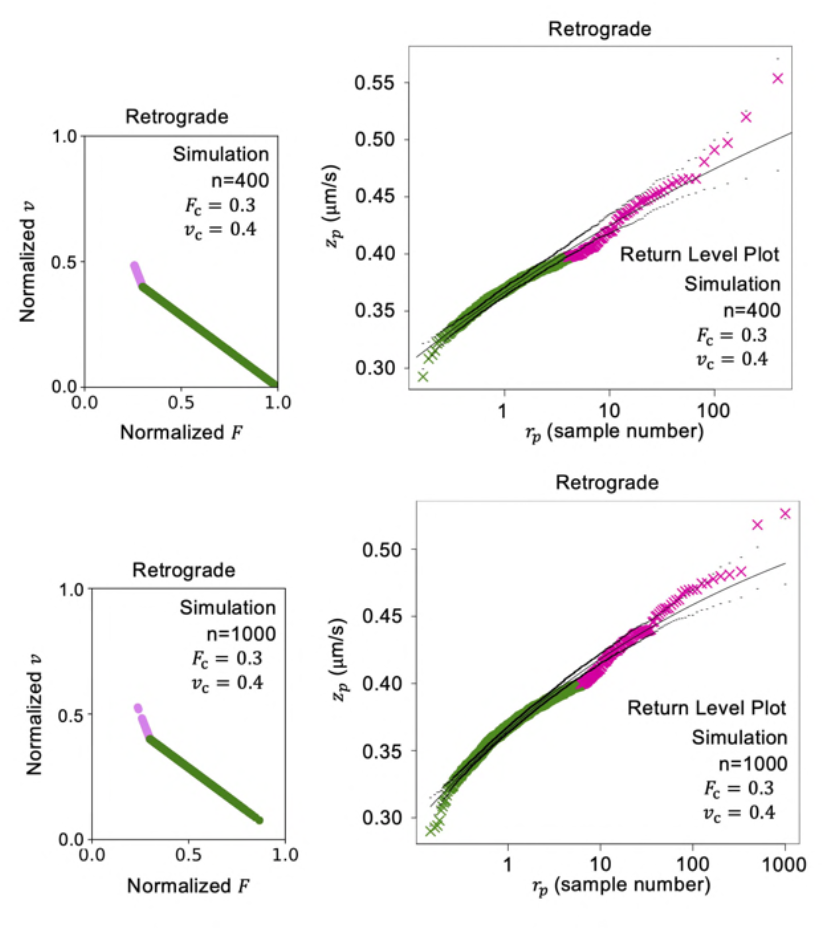
Simulation results of retrograde transport in the cases *n* = 400 and *n*= 1000 for (*F*_c_,*v*_c_) = (0.3,0.4).

**Fig. S4.**
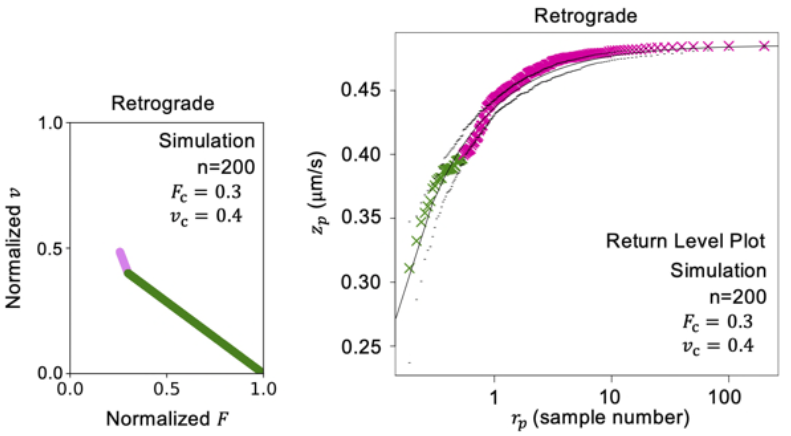
Simulation results of retrograde transport in the case that the cargo size distribution is uniform. (*F*_c_,*v*_c_) = (0.3,0.4).

**Fig. S5.**
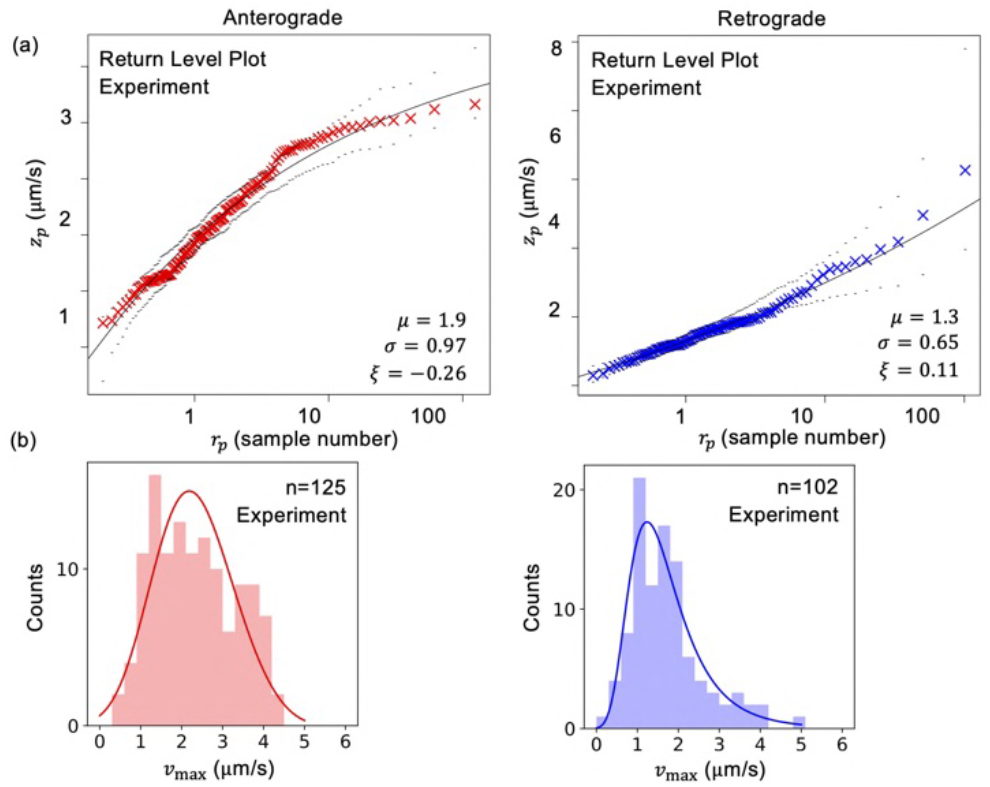
Results of mouse neurons. (a) Return-level plots experimentally measured for anterograde and retrograde transport in the neurons of mouse hippocampal neurons (culture neurons). The time course data was originally obtained by using fluorescence microscopy in the reference [Hayashi *et al*., *Biophys. J*., **120**, 1605-1614 (2021)]. A block is one neuron of the mouse. Each cargo transport was observed and recorded during about 30 sec. One block contains about ten velocity data on average. The black dotted lines, represents the reliable section. (b) Distributions of *v*_max_ for anterograde and retrograde transport.

**Fig. S6.**
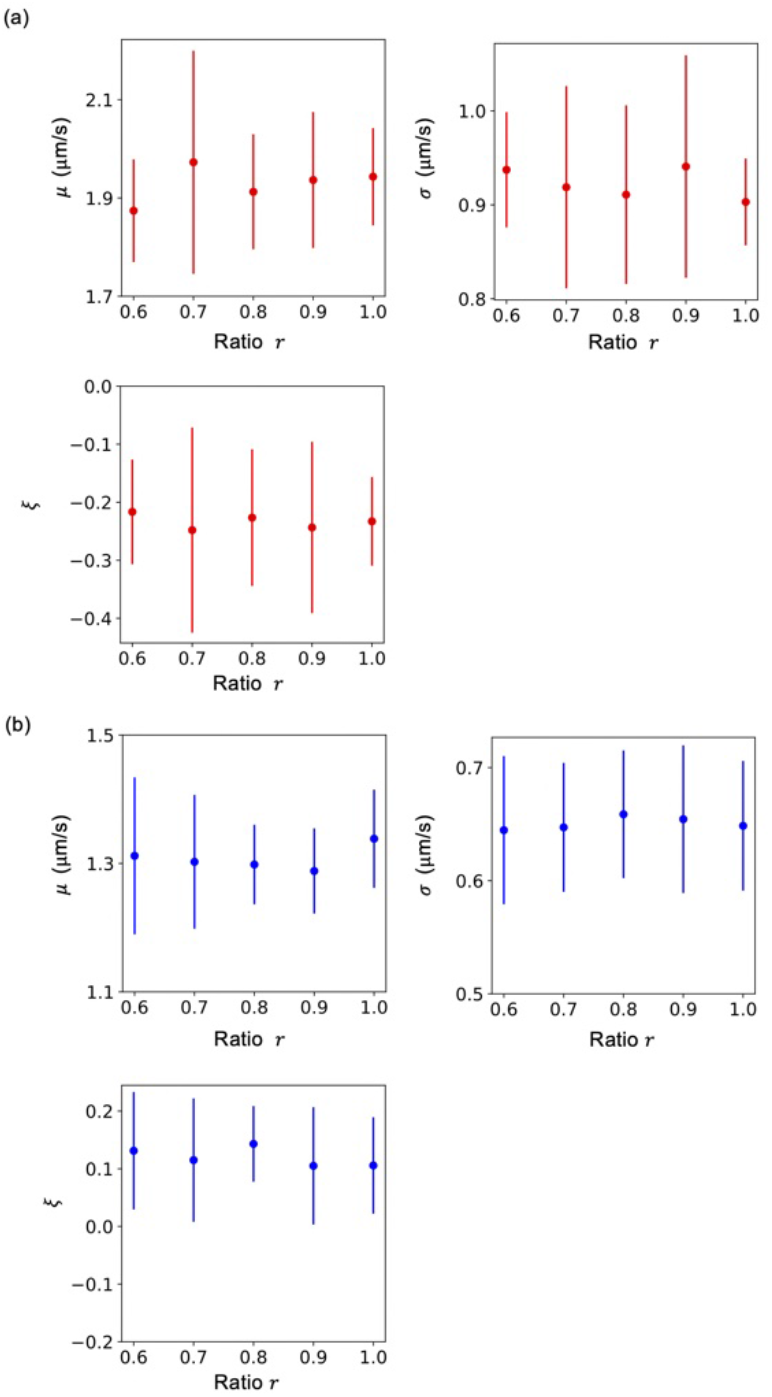
Results of the boot strapping method (mouse). Fitting parameters of *μ, σ*, and *ξ* plotted as a function of ratio (*r*). The bootstrapping analysis was repeated 10 times for each *r*. The error-bars represent the standard deviations (SD).

